# Alternative splicing generates a Ribosomal Protein S24 isoform induced by neuroinflammation and neurodegeneration

**DOI:** 10.1101/2025.03.28.645676

**Authors:** Srivathsa S Magadi, Maria Jonson, Joseph Agi Maqdissi, Lech Kaczmarczyk, Jente J. Zijlstra, Matthew Perkins, Gesine Paul, Martin Hallbeck, Martin Ingelsson, Joel C. Watts, Nicole Reichenbach, Gabor C. Petzold, Pablo B. Lucena, Michael T. Heneka, Walker S. Jackson

## Abstract

Neuroinflammation, particularly that involving reactive microglia, the brain’s resident immune cells, is implicated in the pathogenesis of major neurodegenerative diseases. However, early markers of this process are in high demand. Multiple studies have reported changes in ribosomal protein (RP) expression during neurodegeneration, but the significance of these changes remains unclear. Ribosomes are evolutionarily conserved protein synthesizing machines, and although commonly viewed as invariant, accumulating evidence suggest functional ribosome specialization through variation in their protein composition. By analyzing cell type-specific translating mRNAs from mouse brains, we identify distinct RP expression patterns between neurons, astrocytes, and microglia, including neuron-specific RPs, *Rpl13a* and *Rps10*. We also observed complex expression relationships between RP paralogs and their canonical counterparts, suggesting regulated mechanisms for generating heterogeneous ribosomes. Analysis across brain regions revealed that *Rplp0* and *Rpl13a*, commonly used normalization references, show heterogeneous expression, raising important methodological considerations for gene expression studies. Importantly, we show that *Rps24*, an essential ribosome component that undergoes alternative splicing to produce protein variants with different C-termini, exhibits striking cell type-specific isoform expression in brain. The *Rps24c* isoform is predominantly expressed in microglia and is increased by neuroinflammation caused by aging, neurodegeneration, or inflammatory chemicals. We verify increased expression of S24-PKE, the protein variant encoded by *Rps24c*, in brains with Alzheimer’s disease, Parkinson’s disease, and Huntington’s disease, and relevant mouse models, using isoform-specific antibodies. These findings establish heterogeneous RP expression as a feature of brain cell types and identify *Rps24c*/S24-PKE as a novel marker for neuroinflammation and neurodegeneration.

## Introduction

Neurodegenerative diseases (NDs) such as Alzheimer’s disease (AD), Parkinson’s disease (PD), and Huntington’s disease (HD) are characterized by protein aggregation and disrupted protein homeostasis. These conditions trigger neuroinflammatory responses involving reactive astrocytes and microglia which, while initially protective, can become damaging when chronically activated ^1–9^. The distinct clinical manifestations of each ND reflect differential vulnerability of brain regions and cell types to protein aggregation pathology ^10–23^.

Protein synthesis regulation is fundamental to cellular function, particularly in the brain where distinct cell types maintain unique proteomes and require local translation at sites distal from their cell bodies ^24–27^. Translation of information carried by transcripts is performed by ribosomes, comprising four ribosomal RNAs and approximately 80 ribosomal proteins (RPs). The concept of “specialized ribosomes” has emerged from observations that ribosomal composition can vary between tissues and conditions to selectively translate specific mRNA subsets ^28–34^. While transcriptional control in brain cells is well understood, the contribution of translational regulation through ribosome specialization remains largely unexplored. This specialization could be particularly relevant in the brain, where precise spatial and temporal control of protein synthesis is essential for maintaining neuronal networks, astrocytic support, and microglial responses to pathological stimuli. The specialized ribosome hypothesis requires that subsets of mRNAs contain distinguishing motifs and that these mRNA subsets are selectively translated by ribosomes with unique compositions which could be accommodated with heterogeneous RP expression patterns ^35^.

Ribosomes consist of large and small subunits, and their constituent proteins are named accordingly with L or S prefixes followed by a number. When referring to RP genes or transcripts, the names are italicized, include “*RP”*, with all letters uppercase in human or first letter only uppercase in mouse. Among RPs, S24, encoded by *RPS24* in humans and *Rps24* in mice, is unique as its transcripts undergo alternative splicing to produce protein variants with different C-terminal sequences that are differentially expressed across tissues ^36–42^. *RPS24* expression changes have been associated with various cancers ^42–49^, and mutations in *RPS24* are linked to Diamond-Blackfan Anemia ^50,51^. S24’s location near the mRNA entry tunnel of the ribosome ^52^ suggests a potential involvement in transcript selection.

Here, we report our findings of heterogeneous expression of RPs across major brain cell populations. We identified several neuron-specific RPs and demonstrate distinct patterns of *Rps24* isoform expression between neurons, astrocytes, and microglia. Notably, we find that one isoform, *Rps24c*, is predominantly expressed in microglia and is increased by neuroinflammation. Using newly developed isoform-specific antibodies, we show that the corresponding protein, S24-PKE, serves as a novel marker for neuroinflammation across multiple NDs in both mouse models and patients. This work provides new tools for monitoring neuroinflammatory responses in NDs, and its findings indicate that RPs are heterogeneously expressed across brain cell types which may contribute to cell type-specific translation regulation via specialized ribosomes.

## Results

### Heterogeneous expression of RPs across brain cell types

To investigate potential heterogeneity in RP expression across brain cell populations, we implemented a cell type-specific ribosome-bound mRNA isolation approach using RiboTag knock-in mice ^53^ in which ribosome protein L22 is engineered to express an HA-epitope-tagged variant upon Cre recombinase activation. We employed Cre driver lines specific for microglia (Cx3Cr1-CreERT2) ^54^ and astrocytes (Slc1a3-CreERT2) ^55^ to enable capture of cell type-specific translatome profiles (Supplementary Fig. 1a) across three broad brain regions: front (cortex and striatum), middle (cortex, hippocampus, thalamus, and hypothalamus), and rear (cerebellum, midbrain, and brainstem) (Supplementary Fig. 1b) ^56^. Although we recently found that *Slc1a3* is also expressed in microglia ^57^, we previously verified that the Slc1a3-CreERT2 mouse line drives expression of RiboTag specifically in astrocytes ^55^. Analysis of total mRNAs for region-specific markers confirmed consistent separation of regions (Supplementary Fig. 1b). The specificity of our preparations was validated by robust enrichment of established cell-type markers in RiboTag-captured mRNAs (Fig. 1a). Astrocyte-specific RiboTag translatomes showed significant enrichment of *Apoe*, *Aqp4*, *Fabp7*, *Gja1*, *Gjb6*, and *Slc1a3* across all brain regions ^55,58–60^, while microglia-specific translatomes exhibited high expression of *Aif1*, *Cx3cr1*, *Fcer1g*, *Itgam*, and *Lyz2* ^61,62^.

**Figure 1.**
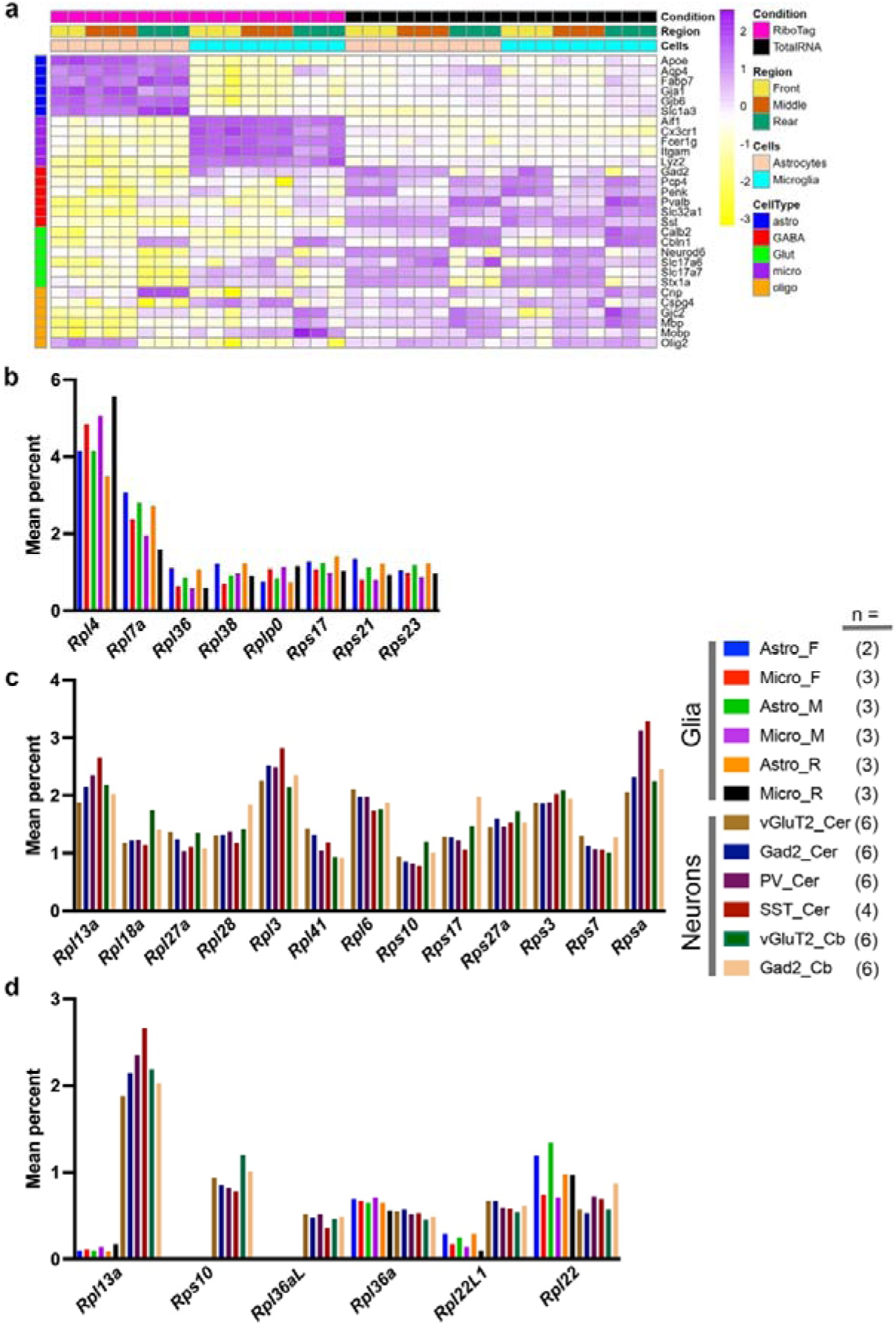
Cell type-specific expression of ribosomal protein transcripts in mouse brain. (a) Expression profiles of cell type-specific markers across brain regions. Heatmap shows scaled expression values from RiboTag and total RNA samples, organized by brain region (Front, Middle, Rear) and cell type (Astrocytes, Microglia). The purple to yellow scale represents Z-scores. (b) Relative abundance of selected ribosomal proteins in astrocytes and microglia. Values represent mean percentage of total RP transcripts. The key includes the number of biologically independent samples per condition in parentheses. (c) Comparison of RP expression across neuronal subtypes. Data shown for vGluT2+, Gad2+, PV+, and SST+ neurons from cerebrum (Cer) or cerebellum (Cb). Values represent mean percentage of total RP transcripts. (d) Expression profiles of neuron specific RPs (*Rpl13a*, *Rps10*, *Rpl36aL* and *Rpl22L1*) and RP paralogs (*Rpl36aL*, *Rpl36a*, *Rpl22L1*, and *Rpl22*) across neurons and glia. Bar heights indicate mean percentage of total RP transcripts.

Different cell types are expected to have varying rates of protein synthesis and thus different amounts of ribosomes. However, if all ribosomes were identical, the ratios of RPs should remain constant across cell types. Therefore, to identify cell type-specific differences in RP expression, we analyzed the relative contribution of each RP transcript to the total RP pool captured by RiboTag. This analysis revealed several RPs with significant differential expression between cell types. For example, *Rpl4* showed higher relative expression in microglia compared to astrocytes across all brain regions, while *Rpl7a* displayed the opposite pattern (Fig. 1b, statistical values in table S1). Furthermore, *Rplp0* and *Rps23* showed elevated expression in microglia, whereas *Rpl36*, *Rpl38*, *Rps17*, and *Rps21* were more abundant in astrocytes in two of the three brain regions examined (Fig. 1b).

To determine whether RP heterogeneity extends to neuronal populations, we reanalyzed our previously published RiboTag data ^63,64^ focusing on non-diseased samples, now comparing three pairs of non-overlapping neuron types: glutamatergic (vGluT2) versus GABAergic (Gad2) neurons from cerebellum and cerebrum (the remainder after removal of the cerebellum, without olfactory bulb), and two GABAergic subtypes - parvalbumin (PV) and somatostatin (SST) neurons from the cerebrum. This analysis revealed neuron type-specific RP expression patterns, with *Rpsa* showing higher expression in PV and SST neurons compared to both Gad2 and vGluT2 neurons from the same region (Fig. 1c, statistical values in table S1). In the cerebellum, *Rps17* and *Rpl28* were enriched in Gad2 neurons relative to vGluT2 neurons, while *Rpl18a* and *Rps10* showed the opposite pattern.

We next wondered if there are differences in RP expression between neurons and glia. We recognized an important difference between the studies was that the regions studied were not identical; the neuron study examined the cerebellum and cerebrum which are roughly comparable to the rear sections and the front and middle sections of the glial study, respectively. Nonetheless, all neuron and glia samples (i.e., previous and current data sets) were prepared from nine-month-old mice with the same genetic background (129S4) and were sequenced together on the same flow cell. This analysis uncovered striking differences between neurons and glia, where *Rpl13a*, *Rps10*, *Rpl36aL*, and *Rpl22L1* showed substantially higher expression in neurons compared to glial cells (Fig. 1d). Notably, the latter two are paralogs of canonical RPs *Rpl36a* and *Rpl22*. We observed that *Rpl22L1*, which increases expression in response to reduced expression of *Rpl22* ^65^, showed an inverse relationship with *Rpl22* in astrocytes, but no such reciprocal pattern was detected between *Rpl36aL* and *Rpl36a*.

To verify heterogeneous RP expression at the protein level, we performed immunofluorescence analysis of ribosomal protein S10 in cortical brain sections. Co-labeling with cell type-specific markers revealed that while most neurons expressed S10, GFAP-positive astrocytes showed negligible expression (Fig. 2). Although antibody compatibility issues prevented assessment of S10 with microglial marker expression, S10 was not observed in cells sized or shaped like microglia. This cellular distribution pattern provides independent validation of our RiboTag-based findings and demonstrates that RP heterogeneity exists at both transcript and protein levels.

**Figure 2.**
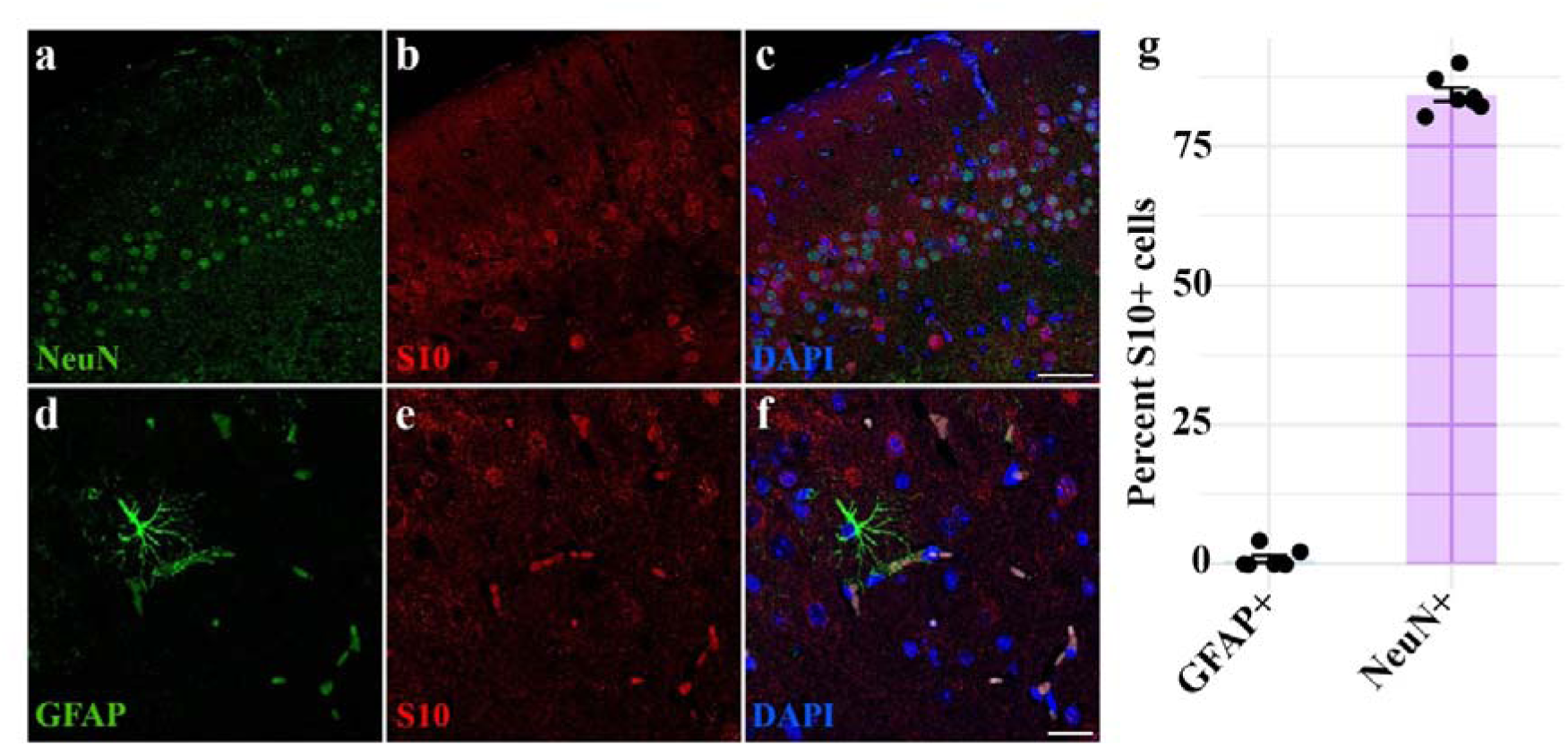
Neuron-specific expression of ribosomal protein S10 in mouse cortex. (a-c) Immunofluorescence image of S10 in cortical neurons. Representative images showing NeuN (green), S10 (red), and DAPI (blue). Scale bar, 50 µm; n = 3 mice. (d-f) S10 expression in relation to astrocytes. GFAP (green), S10 (red), and DAPI (blue). Scale bar, 50 µm; n = 3 mice. (g) Classification of S10-positive cells. Data show percentage of S10+ cells colocalizing with NeuN or GFAP. Individual points represent independent fields of view; bars show mean values; n = 3 mice, 10 fields per mouse.

### Cell type-specific expression of *Rps24* isoforms in healthy and diseased brain

Having identified multiple candidates for heterogeneous RP expression across brain cell types, we investigated whether alternative mRNA splicing could provide another pathway for producing ribosome diversity. We focused on *Rps24*, unique among RP genes for its tissue-specific alternative splicing that produces protein variants with different C-terminal ends ^38^. Remarkably, *Rps24* is one of the most differentially alternatively spliced mRNAs across tissues ^40,41^. Furthermore, these splicing patterns are conserved between humans and mice ^38^ and the protein sequences are identical in human and mouse.

Due to inconsistencies in isoform nomenclature, we devised a systematic naming scheme where specific isoforms in mouse and human share the same designation: transcript isoforms named as *Rps24a, Rps24b, and Rps24c* encode protein isoforms S24-3K, S24-2K, and S24-PKE, respectively (Fig. 3a and Supplementary Discussion). We created custom-designed droplet digital PCR (ddPCR) assays and verified their quantitative accuracy with recombinant RNAs, (Fig. 3a, Supplementary Fig. 2,). Using these assays, we quantified the three major splice forms across tissues and found distinct tissue-specific distributions, with *Rps24a* predominating in heart, *Rps24b* in brain, and *Rps24c* in blood, kidney, liver, lung, and spleen (Fig. 3b, statistical values in table S2). Notably, mouse and human brain showed remarkably similar isoform ratios – *Rps24a*: 11.8% vs. 13.5%, *Rps24b*: 85.4% vs. 85.3%, and *Rps24c*: 2.8% vs. 1.2%, respectively. These results are consistent with previous RNAseq analyses ^38^, indicating that RNAseq can also accurately quantify *Rps24* splice form ratios.

**Figure 3.**
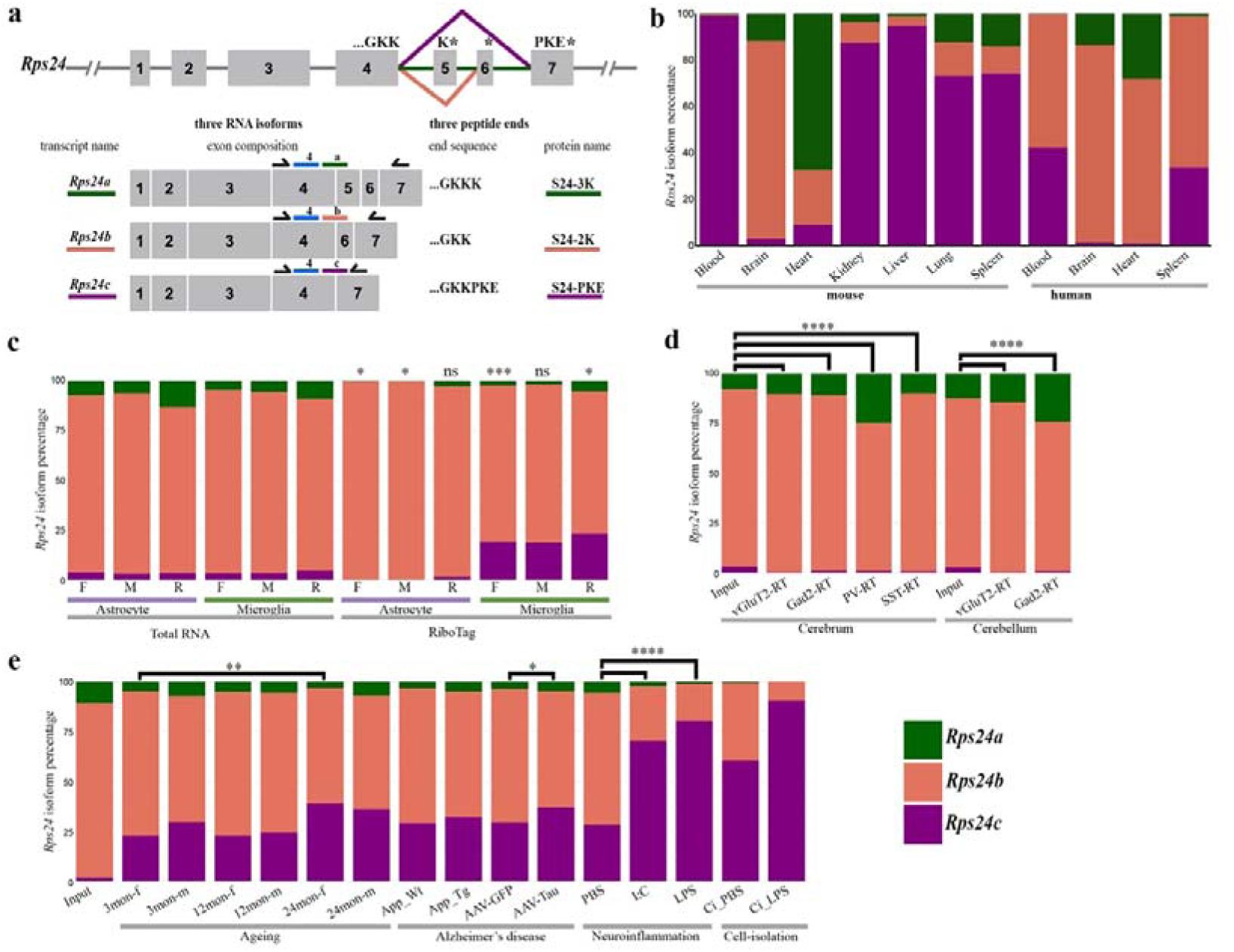
Differential expression patterns of *Rps24* isoforms. (a) Top, the gene structure and alternative splicing pattern of *Rps24*. Exons are depicted as grey boxes and key amino acids and stop codons (*) are above them. Below, RNA isoforms (*Rps24a*, *b*, *c*), and resulting protein variants (S24-3K, 2K, PKE). (b) Tissue-specific distribution of *Rps24* isoforms in mouse and human. Stacked bars show relative abundance of *Rps24a* (green), *Rps24b* (salmon), and *Rps24c* (purple); n = 3 independent samples per tissue. (c) Brain region-specific *Rps24* isoform expression in glia. Statistical comparisons of *Rps24c* levels are made between RiboTag samples and their corresponding Total RNA samples from three brain regions (F, M, R). *P < 0.05, **P < 0.01; one-way ANOVA with Tukey’s post-hoc test; replicate numbers are shown in Fig 1. (d) Neuronal subtype-specific *Rps24* profiles from RiboTag. Statistical comparisons of *Rps24c* levels are made between RiboTag samples and their corresponding Total RNA samples. ****P < 0.0001; two-tailed Student’s t-test; replicate numbers are shown in Fig 1. (e) Dynamic regulation of *Rps24* isoforms in disease models and inflammation. Statistical comparisons of *Rps24c* levels are made between RiboTag samples. Analysis includes aging (3-24 months), AD models, and inflammatory conditions. *P < 0.05, **P < 0.01, ****P < 0.0001; one-way ANOVA with Tukey’s post-hoc test; n = 3-4 mice per condition.

Cell type-specific RiboTag analysis revealed striking differences in *Rps24* isoform expression between brain cell populations. While both astrocytes and microglia showed low *Rps24a* levels (0-2.9% and 1.8-5.3%, respectively), they differed markedly in *Rps24b* and *Rps24c* expression. Astrocytes expressed predominantly *Rps24b* (95.5-100%) with minimal *Rps24c* (0-1.6%), whereas microglia showed lower *Rps24b* (71.5-79.4%) and substantially higher *Rps24c* levels (18.7-23.2%; Fig. 3c, statistical values in table S2). Neuronal populations also displayed distinct patterns, with higher *Rps24a* expression (9.7-24.6%) than glial cells and low *Rps24c* levels (0.2-1.0%) similar to astrocytes but distinct from microglia. Regional differences were also evident, particularly in GABAergic neurons, which expressed 10.7% *Rps24a* in the cerebrum versus 24.0% in the cerebellum (Fig. 3d).

The elevated microglial expression of *Rps24c* prompted us to examine data from a study investigating RiboTag-captured mRNAs from microglia under various inflammatory conditions ^66^. During aging, microglial *Rps24c* levels increased from 22.7-29.5% at 3 months to 36.0-39.0% at 24 months, with sex-specific variations (Fig. 3e). In a transgenic mouse model of AD-related amyloid-β brain pathology, 9-month-old APP/PS1 mice ^67^ showed no significant change in *Rps24c* levels, but a virus-based model of Tauopathy showed an increase from 29.3% to 37.0% (Fig. 3e). Acute inflammatory challenges induced even more dramatic changes. Treatment with polyinosinic:polycytidylic acid (I:C) or lipopolysaccharide (LPS) increased microglial *Rps24c* levels from 28.1% to 70.3% and 80.4%, respectively. Moreover, physical isolation of microglia by flow cytometry elevated *Rps24c* to 60.5%, which further increased to 90.4% upon LPS stimulation (Fig. 3e). These changes occurred alongside overall increases in RP expression (Supplementary Fig. 3), suggesting broader translational remodeling during microglial activation.

### S24-PKE expression correlates with pathological features in NDs

Having established that *Rps24c* is induced in microglia during aging and neuroinflammation, both associated with ND, we asked if these changes were reflected at the protein level. To detect the corresponding protein isoform, S24-PKE, we generated specific antibodies by immunizing mice and rabbits with peptides containing the unique C-terminal sequence. Through systematic screening, we identified hybridoma clones and sera that specifically bound S24 peptides ending with -PKE while showing no reactivity to those ending with -KK or -KKK. We validated antibody specificity using western blots of cell lysates expressing V5-tagged S24 isoform variants (Fig. 4a). Initial probing with V5 antibody confirmed expression of all three S24 variants and a V5-GFP control (Fig. 4b). Both mouse and rabbit anti-S24-PKE antibodies exclusively detected V5-S24-PKE but not V5-S24-2K or V5-S24-3K. As expected, the antibodies also recognized endogenous S24-PKE, which migrated at a lower molecular weight than the V5-tagged variants.

**Figure 4.**
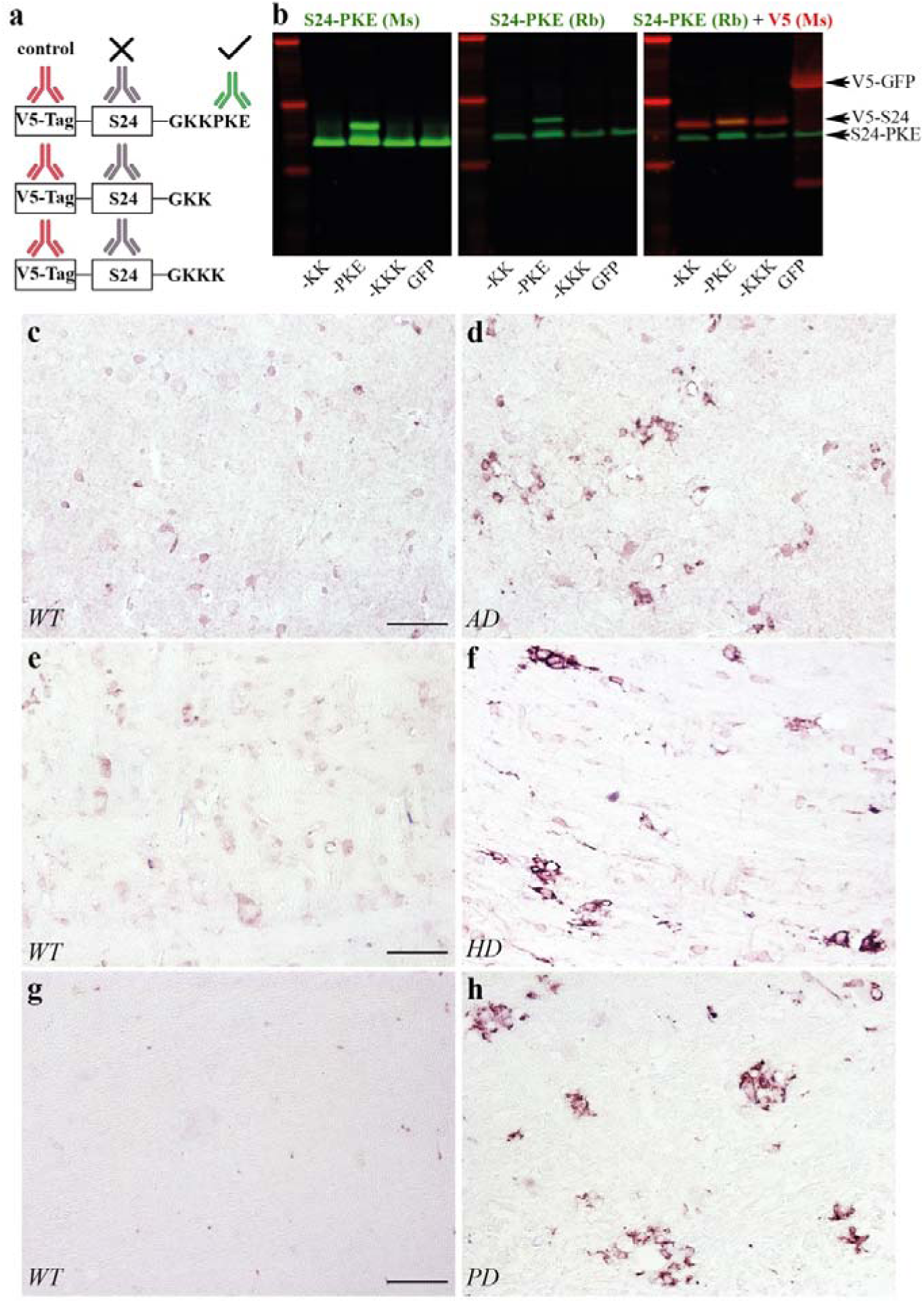
Validation and characterization of S24-PKE expression in neurodegenerative disease mouse models. (a) Design of V5-tagged S24 constructs for antibody validation. (b) Western blot validation of S24-PKE antibodies using lysates from HEK cells expressing the constructs depicted in A. The left and middle strips were duplicates made from the same gel and lysates. Representative blots from three independent experiments. (c-h) S24-PKE immunoreactivity in mouse models of neurodegeneration. Cortical sections from AD model (c, d), brainstem sections from HD model (e, f), and brainstem sections from PD model (g, h). Scale bars, 50 µm; n = 3 mice per condition.

Using these validated antibodies, we examined formalin-fixed paraffin-embedded (FFPE) brain sections from multiple mouse models of ND. In APP/PS1 mice, we observed S24-PKE labeling in cortex and hippocampus of mutant mice, while control littermates showed minimal staining (Fig. 4c, d). In the HdhQ200 HD model mice ^68,69^ backcrossed to the 129S4 background ^64^, we detected S24-PKE expression in brainstem and cerebellum at 18 months, with minimal signal in age-matched controls (Fig. 4e, f). In a PD model using M83 transgenic mice expressing A53T mutant α-synuclein ^70^ injected with pathological α-synuclein aggregates ^71^, we observed S24-PKE expression that was largely absent in controls (Fig. 4g, h). Dual immunofluorescence labeling revealed that S24-PKE-positive cells were predominantly Iba1-positive microglia (Fig. 5). Finally, we sought to determine how S24-PKE expression related to protein aggregates. In PD model mice, S24-PKE was found in regions containing high levels of phosphorylated α-synuclein (P-Syn) deposits (Fig. 6a-c). Similarly, in vibratome sections of formaldehyde-fixed APP/PS1 mice (18 mos old), S24-PKE-expression was observed primarily near amyloid plaques in Iba1+ microglia but not in GFAP+ astrocytes (Fig. 6d-h). Therefore, S24-PKE is expressed in distinct mouse models of ND, primarily in microglia, often near sites with protein aggregates.

**Figure 5.**
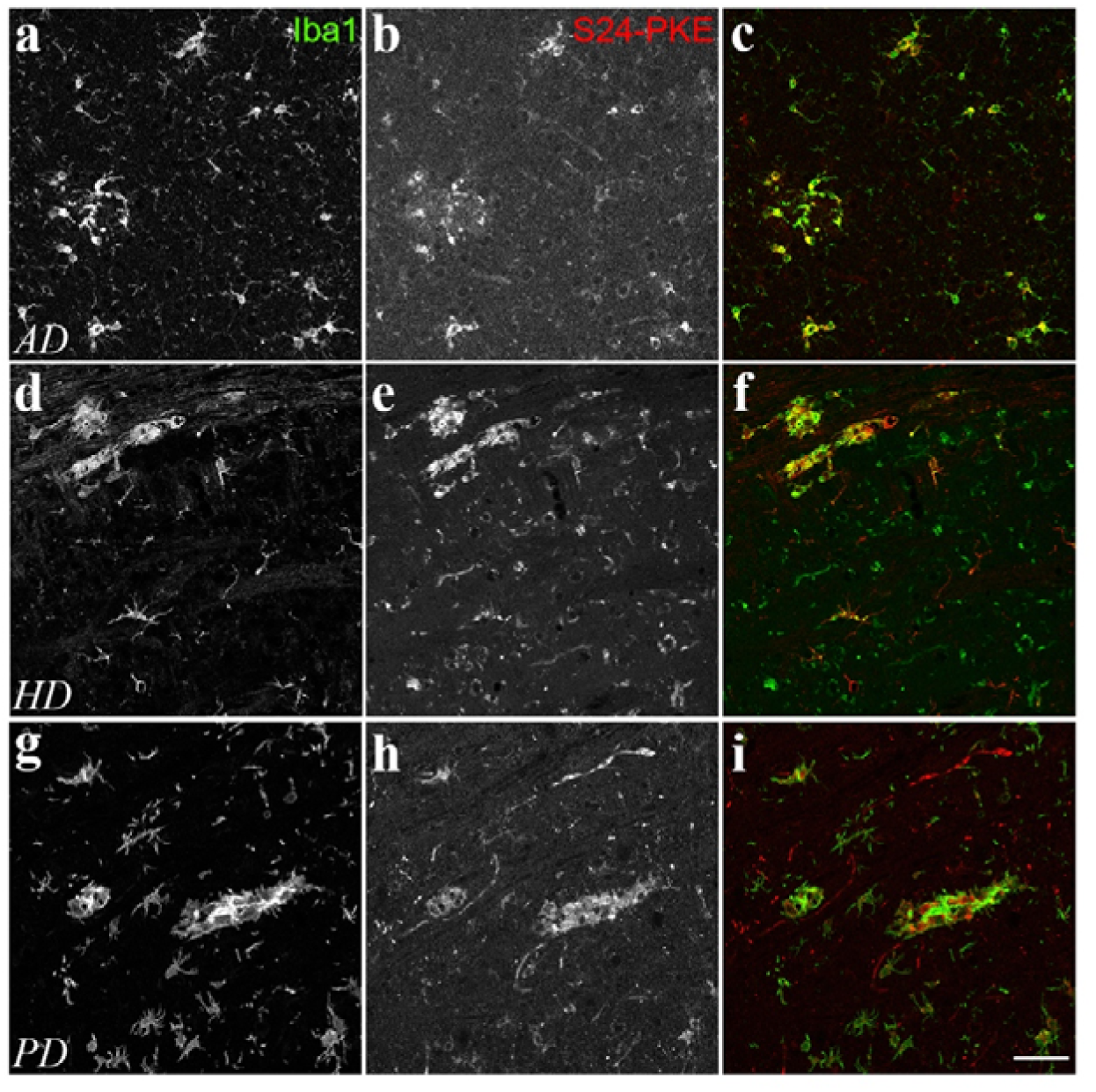
Microglial localization of S24-PKE in neurodegenerative diseases. (a-i) Double immunofluorescence analysis of S24-PKE and Iba1 in FFPE sections from ND models. Representative images from (a-c) AD cortex, (d-f) HD brainstem, and (g-i) PD brainstem. Individual channels shown in grayscale, merge displays colocalization. Scale bars, 50 µm; n = 3 mice per condition.

**Figure 6.**
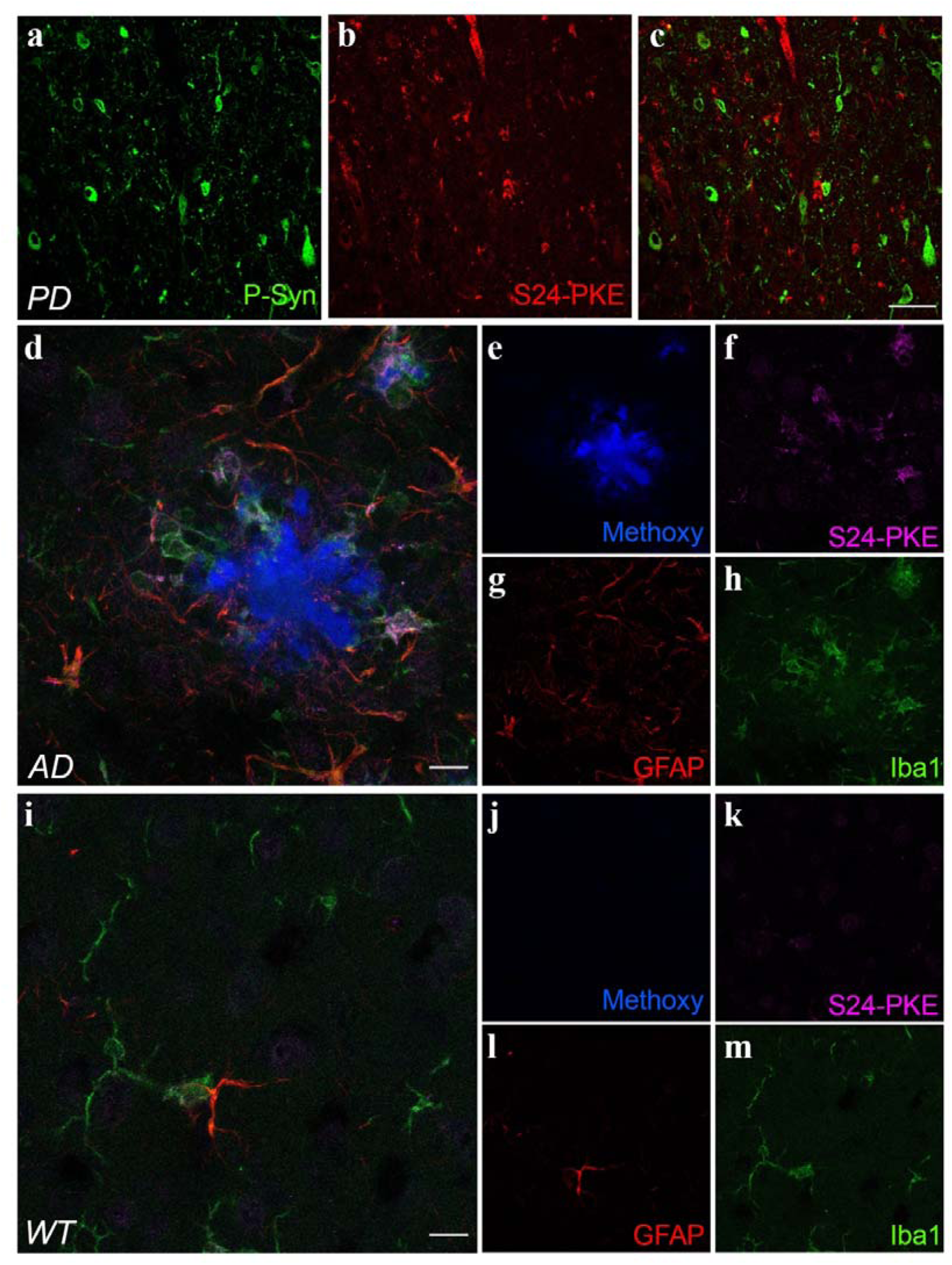
Association of S24-PKE with pathological protein deposits. (a-c) Spatial relationship between S24-PKE and phosphorylated α-synuclein in PD model. Scale bar, 50 µm. (d-h) Multi-label analysis in APP/PS1 cortex showing the relationship between S24-PKE, amyloid plaques, and glial markers. Scale bar, 50 µm. (i-m) Corresponding analysis in wild-type controls. Scale bar, 50 µm. Representative images from n = 3 mice per condition.

Finally, to determine if S24-PKE expression is induced in human brains with ND, we analyzed the labeling of our antibodies on FFPE post-mortem brain tissue from patients with AD, HD, and PD. Control sections consistently showed minimal immunoreactivity. In contrast, all examined HD and PD sections, and two of three AD sections showed S24-PKE staining in cells morphologically consistent with microglia and in certain vascular structures (Fig. 7). Immunofluorescence analysis confirmed S24-PKE expression in Iba1+ cells and in some Iba1-structures with vascular morphology (Fig. 8). These observations indicate that S24-PKE expression is induced in human neurodegenerative conditions, though further studies will be needed to fully characterize its expression pattern and function.

**Figure 7.**
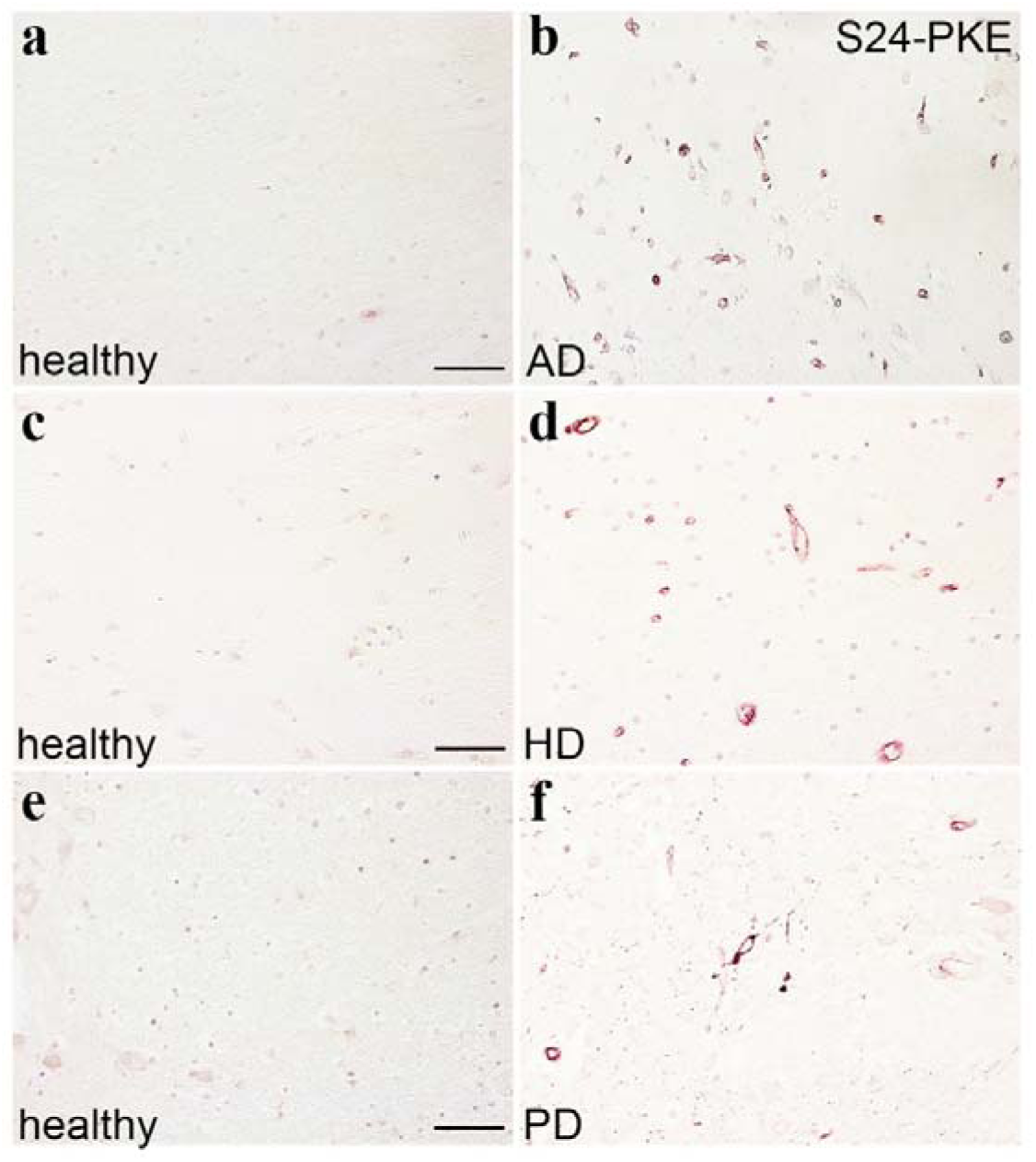
S24-PKE expression in human neurodegenerative diseases. (a-f) Immunohistochemical analysis of S24-PKE in human brain tissue. Representative images from (b) AD cortex, (d) HD striatum, and (f) PD basal ganglia, with age matched controls. Scale bars, 50 µm; n = 2 AD, n = 3 HD, n = 2 PD.

**Figure 8.**
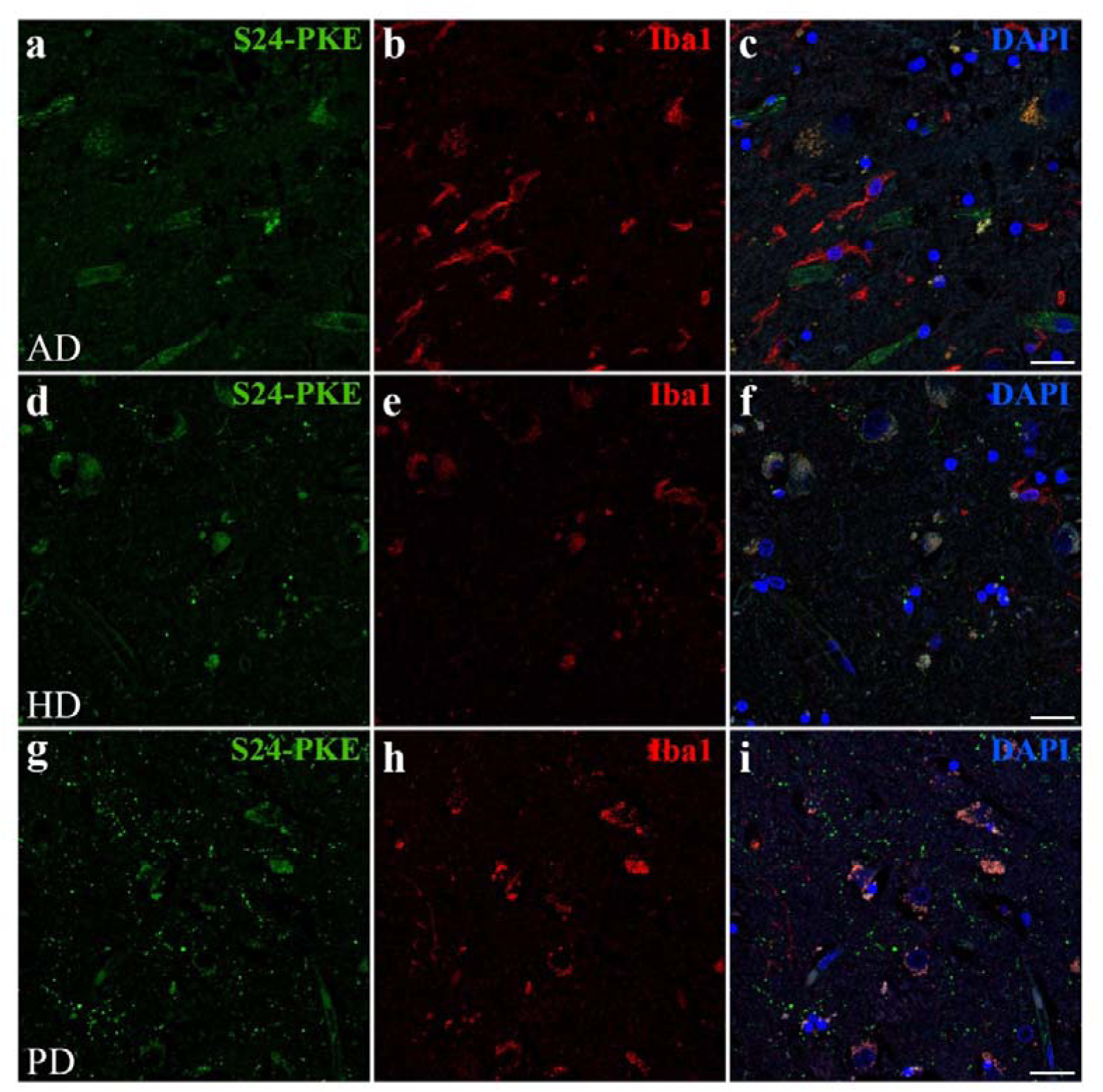
Partial co-localization of S24-PKE and Iba1 in human NDs. (a-i) Triple co-localization analysis showing S24-PKE, Iba1, and DAPI in (a-c) AD cortex, (d-f) HD striatum, and (g-i) PD basal ganglia. Scale bars, 50 µm; n = 2 AD, n = 3 HD, n = 2 PD.

## Discussion

To investigate alterations in protein synthesis during NDs, we recently used the RiboTag method to analyze the cell type-specific translatome of mouse models of four NDs ^60,63,64^. From these and similar studies we noticed that the expression of mRNAs encoding RPs often change ^23^. While these observations might reflect modulation of ribosome biogenesis, only a fraction of RPs change. An alternative explanation is that RP expression is altered to shift ribosome heterogeneity, thereby recalibrating the pool of specialized ribosomes. This knowledge gap led to the current study, which reveals previously unrecognized heterogeneity in ribosomal protein expression across brain cell types and demonstrates dynamic regulation of S24-PKE in neurodegeneration.

### Heterogeneous Expression of RPs Suggests Cell Type-Specific Translation Mechanisms

Despite sharing identical DNA sequences, brain cells develop distinct morphologies and functions through cell type-specific gene expression programs. While transcriptional regulation has been extensively studied, our findings suggest an additional layer of control through specialized ribosomes. We identified distinct patterns of RP expression across brain cell types, including heterogeneous expression of *Rpl36aL* and *Rpl22L1*, paralogs of canonical RPs that can incorporate into ribosomes. These paralogs represent promising candidates for generating specialized ribosomes with distinct translational preferences, similar to S27L, which selectively translates specific transcript pools ^34^, and L39L, which is needed for sperm-specific ribosomes ^72^. Like L22L1, both S27L and L39L show inverse expression relationships with their canonical paralogs ^34,72^. This pattern of paralog switching may represent a broader mechanism for generating specialized ribosomes than previously appreciated.

### *Rps24c*/ S24-PKE as a Novel Marker of Neuroinflammation and NDs

The identical protein sequences and the remarkable conservation of isoform ratios between mouse and human brain (*Rps24a*: 11.8% vs. 13.5%, *Rps24b*: 85.4% vs. 85.3%, *Rps24c*: 2.8% vs. 1.2%, respectively) suggest strong evolutionary pressure to maintain specific proportions. A recent survey of *RPS24* isoforms in tumors derived from many tissues across the body found that *RPS24c* tends to increase ^42^. Interestingly, in brain tumors, *RPS24a* was strongly reduced but *RPS24c* was apparently unchanged ^42^. In our study, while total *Rps24* expression remained relatively constant, we uncovered striking cell type-specific patterns among its isoforms. We found that *Rps24a* was primarily expressed by neurons to variable levels, depending on the neuron type. In contrast, *Rps24c* was primarily expressed by microglia and, strikingly, *Rps24c* expression was further increased by diverse neuroinflammatory stimuli. Importantly, our observations that cell isolation procedures significantly alter *Rps24c* expression coincided with expression of genes associated with reactive microglia ^66^, along with heterogeneous expression of common reference genes like *Rplp0* and *Rpl13a*, raise critical methodological considerations for studying microglia and normalizing gene expression data.

The development of S24-PKE-specific antibodies addressed a significant gap in protein-level analysis ^41^. Notably, S24-PKE expression in the HdhQ200 model preceded typical neuroinflammatory markers ^64^, suggesting its potential as an early indicator of cellular stress. It is important to note that a low level of S24-PKE was detected in brains from aged (at least 16 months old) healthy mice, consistent with increased levels of *Rps24c* during aging, although aged healthy human brains were invariably negative. The conservation of S24-PKE expression patterns between brain tissues from mouse disease models and patients further supports its utility as a disease marker.

### Mechanistic Implications and Future Directions

The biological significance of *Rps24*/S24 isoforms extends beyond their association with cancer ^43,45^ and hypoxia responses ^73,74^. The coincident increase in global RP expression with *Rps24c* induction during aging and inflammation, combined with S24’s role in ribosomal biogenesis ^51^, suggests involvement in broader translational remodeling. This is particularly relevant given the increased RP expression observed in neurodegenerative conditions ^75–77^.

A surprising paradox emerged with our antibodies: despite *Rps24c* association with intact ribosomes, S24-PKE protein levels remain low in healthy brains. While this might reflect regulated translation, evidence suggests a shorter half-life for S24-PKE compared to S24-KK ^73^. Future research requires careful *in vivo* investigation of isoform expression consequences in health and disease. Given the tight regulation of *Rps24* expression and documented gene dosage compensation ^51,78^, studies manipulating the endogenous gene will likely yield more reliable results than random integration transgenes or viral vectors, despite the great effort required. Likewise, the specific association of *Rps24c*/S24-PKE with neuroinflammatory conditions and NDs suggests potential therapeutic applications, though careful consideration of *Rps24*’s tight expression regulation and dosage compensation will be crucial for such approaches. Meanwhile, our S24-PKE antibody provides a valuable tool for investigating brain pathologies associated with NDs or neuroinflammation.

In conclusion, these findings significantly expand on previous evidence of ribosome heterogeneity as a feature of brain regions and cell types ^16,55^ and identify *Rps24c*/S24-PKE as a novel marker for NDs and neuroinflammation, providing new tools for studying neurological diseases while suggesting potential therapeutic approaches by controlling expression of *RPS24*/S24 isoforms.

## Methods

### Ethics and Tissue Samples

Animal experiments were approved by the Landesamt für Natur, Umwelt und Verbraucherschutz Nordrhein-Westfalen (84-02.04.2013.A169, 84-02.04.2013.A128, 84-02.04.2016.A442) and Linköpings djurförsöksetiska nämnd (14741-2019). Human brain samples were obtained from brain banks at Uppsala University, Lund University, and University of Michigan with appropriate ethical approvals and informed consent.

### Mouse Models

RiboTag mice ^53^ were crossed with Slc1a3-CreERT2 ^55^ and Cx3cr1-CreERT2 ^54^ mouse lines on a 129S4 background. Cre activity was induced with Tamoxifen (Sigma-Aldrich, T5648; 100 mg/kg, i.p.) for three consecutive days. Additional models included APP/PS1 ^67^, TgM83 ^70^ injected with pathological aggregates ^71^, and HdhQ200/Q7 mice ^64^. Animals were housed in ventilated cages with 12-hour light/dark cycles at 23 ± 2°C with ad libitum access to water and standard chow.

### RiboTag Analysis

Brain regions were dissected, flash-frozen, and processed for RiboTag immunoprecipitation (Supplementary Fig. 1a) ^60^. Tissue was homogenized in polysome buffer (10 mM HEPES pH 7.4, 150 mM KCl, 5 mM MgCl2, 0.5 mM DTT, 100 μg/mL cycloheximide) with RNase inhibitors. Lysates were incubated with anti-HA antibody (Roche, 11583816001) and Protein G magnetic beads (Thermo Fisher Scientific, 10004D). RNA was extracted using RNeasy Mini Kit (Qiagen, 74104).

### RNA Sequencing

Libraries were prepared using TruSeq Stranded mRNA Library Prep Kit (Illumina, 20020594) and sequenced on NovaSeq 6000 with 150 bp paired-end reads at National Genomic Infrastructure (NGI), Sweden. Reads were aligned to mouse genome (mm10) using STAR aligner (v2.7.3a). For RP expression analysis, the number of reads mapping to RP genes was summed and used to calculate the percent of RP reads mapping to each RP gene

### *Rps24* Isoform ddPCR Analysis

Total RNA was extracted from mouse tissues (blood, brain, heart, lung, kidney, and spleen) using the Norgen Total RNA Purification Kit (Norgen Biotek, 17200) according to manufacturer’s instructions. RNA was eluted in RNase-free water and quantified using NanoDrop 2000 Spectrophotometer (Thermo Fisher Scientific, ND-2000). Human blood samples were processed similarly, while human brain, heart, and spleen RNA samples were obtained commercially (Thermo Fisher Scientific; Brain: AM7962; Heart: AM7966; Spleen: AM7970). RNA samples were stored at −80°C until use. cDNA was synthesized using the Protoscript II Reverse Transcriptase (NEB, M0368L) with random hexamer and oligo dT primers. For each reaction, 500 ng total RNA was first denatured at 65°C for 5 minutes, followed by reverse transcription at 42°C for 50 minutes, and enzyme inactivation at 80°C for 5 minutes according to manufacturer’s protocol. The resulting cDNA was diluted 1:5 in nuclease-free water and stored at −20°C until further use. *Rps24* isoforms were quantified using droplet digital PCR (ddPCR). Forward primers were designed to target the shared exon 4, while reverse primers targeted exon 7. Isoform-specific sequences were: *Rps24a*: 5’-TGGCAAAAAGAAATGAAGTG-3’ *Rps24b*: 5’-TGGCAAAAAGTGAGCTGGAG-3’ *Rps24c*: 5’-TGGCAAAAAGCCGAAGGAGT-3’ Common *exon 4*: 5’-AATGTTGGTGCTGGCAAAAA-3’. Custom probes targeting isoform-specific junctions were labeled with either FAM or HEX fluorescent dyes: *Rps24a* (exon 4-5, HEX), *Rps24b* (exon 4-6, HEX), and *Rps24c* (exon 4-7, FAM). A common exon probe targeting exon 4 (not overlapping with isoform-specific regions) was labeled with either HEX or FAM. Probe specificity was validated using plasmids containing isoform-specific sequences from both human and mouse (Supplementary Fig. 2a). ddPCR reactions contained 10 μL QX200 ddPCR EvaGreen Supermix (Bio-Rad, 1864034), primers (900 nM final), probes (250 nM final), and 5 μL cDNA template (equivalent to 0.2 ng/µL RNA) in 20 μL total volume. Analysis was performed using QX200 Droplet Reader and QuantaSoft software (Bio-Rad). Isoform-specific droplet populations are shown in Supplementary Fig. 2b.

### Cell Culture and Antibody Validation

HEK293T cells were maintained in DMEM with 10% FBS and transfected with V5-tagged Rps24 isoforms using Lipofectamine 2000 (Thermo Fisher, 11668019). Lysates were analyzed by western blot using mouse S24-PKE (1:2000), rabbit S24-PKE (1:500), and mouse-anti-V5 (1:5000, Invitrogen, R96025) antibodies. Secondary antibodies included donkey-anti-rabbit/mouse IRDye 800CW (1:20,000, Li-Cor, 926-32213/926-32212).

### Immunohistochemistry

Brain sections (4 μm) were deparaffinized in xylene, rehydrated through graded ethanol solutions (100-50%) and PBS, with 5-minutes per step. Epitope retrieval was performed in 0.01M citrate buffer (pH 8.0) by steaming for 20 minutes, followed by cooling at room temperature for 15 minutes. For chromogenic detection, sections were treated with 0.3% H2O2, blocked with 2.5% normal horse serum, and incubated with mouse S24-PKE antibody (3.4 μg/ml). Detection used ImmPRESS HRP polymer kit (Vector Laboratories, MP-7401) and ImmPACT VIP substrate (Vector Laboratories, SK-4605).

### Immunofluorescence

Brain sections were processed for deparaffinization and antigen retrieval as described for immunohistochemistry. Autofluorescence was quenched using TrueBlack Lipofuscin Autofluorescence Quencher (Biotium, 23007). Sections were blocked in 2.5% normal horse serum (PBS) for 30 min before overnight primary antibody incubation at 4°C. Primary antibodies: S24-PKE-1 (3.4 μg/ml), Iba1 (1:200, Wako, 019-19741), Rps10 (GeneTex, GTX101836), anti-phospho α-synuclein (1:200, Wako, 015-25191) and GFAP (1:2000, Millipore, MAB360). Secondary antibodies (all 1:500): Alexa Flour 488, Alexa Fluor 647, and Cy3 (Jackson Immunoresearch). Sections were mounted using Vectashield vibrance antifade mounting media (Vector Laboratories, H-1700) and imaged using a Zeiss LSM 800 confocal microscope with 20x objective.

For amyloid plaque visualization, brains were perfused with ice-cold PBS for 2 min and fixed in 4% PFA–PBS (24h, 4°C). Free-floating sagittal sections (40 μm) were generated using a Leica VT1000 S vibratome. Sections were permeabilized in PBS-0.5% Triton X-100 and underwent citrate buffer antigen retrieval (0.01M citrate, pH 6.0, 0.05% Tween-20) by microwave treatment. After blocking (1% BSA in PBS-Triton), sections were incubated overnight at 4°C with primary antibodies: GFAP (1 μg/ml, ThermoFisher Scientific, 2.2B10), IBA-1 (0.5 μg/ml, Abcam, 5076), and mouse S24-PKE (3.7 μg/ml). Secondary antibodies (Invitrogen): AlexaFluor-488, AlexaFluor-568, and AlexaFluor-647. Autofluorescence was reduced using Sudan Black solution (0.1% in 70% ethanol). Nuclei were visualized with DAPI (1:5000) or Methoxy-X04 (10 μM in ethanol). Sections were mounted using Fluoromount-G™ (ThermoFisher, 00-4958-02) and imaged using a Leica SP8 microscope.

### Statistical Analysis

Data were analyzed using R (v4.0.3). Comparisons between groups used Student’s t-test or Mann-Whitney U test as appropriate. Multiple comparisons used one-way ANOVA with Tukey’s post-hoc test or Kruskal-Wallis with Dunn’s post-hoc test. P < 0.05 was considered significant. Data are presented as mean ± SEM unless otherwise stated.

## Data Availability

RNA-seq data generated in this study have been deposited in the NCBI Gene Expression Omnibus (GEO) under accession number (GSE289868). Additional datasets analyzed in this study include previously published RiboTag data from neuronal subtypes (GSE198063; Bauer et al., 2022, GSE199837 Bauer et al., 2023) and microglia under various inflammatory conditions (GSE117646; Kang et al., 2018). Additional data available from corresponding author upon reasonable request.

## Acknowledgements

We thank Drs. Maria Ntzouni and Vessa Loitto for assistance with histology and microscopy. Animal care was provided by the Core Facility for Laboratory Animals (CBR). Computational support was provided by the National Genomics Infrastructure Sweden and National Bioinformatics Infrastructure Sweden. Analyses were performed using the NextFlow Core pipeline. This work was supported by grants from the Michigan Brain Bank (P30AG053760/P30AG072931 University of Michigan Alzheimer’s Disease Core Center) to M.P., the Wallenberg Center for Molecular Medicine to W.S.J., Konung Gustaf V:s och Drottning Victorias Stiftelse to W.S.J., Hereditary Disease Foundation to S.M., Lions Forskningsfond to M.J., Parkinsons Stiftelse to W.S.J., LiU Systems Neurobiology to W.S.J., Hjärnfonden to M.J., and Fonds National de la Recherche PEARL program (FNR/16745220) to M.T.H.

## Author Contributions

S.S.M. and W.S.J. developed the concept of the project, S.S.M., M.J., J.A.M., L.K., J.Z., and P.B.L. performed experiments, M.P., G.P., M.H., M.I., J.C.W., N.R., and G.C.P. provided key materials, S.S.M., M.J., M.T.H., and W.S.J. obtained funding, and M.T.H., and W.S.J. supervised research. S.S.M. and W.S.J. wrote the original draft and all authors edited the manuscript.

## Additional Information

**Supplementary information** The online version contains supplementary material

**Correspondence** and requests for materials and code should be addressed to W.S.J.

