## Supplementary material for "Alternative splicing generates a Ribosomal Protein S24 isoform induced by neuroinflammation and neurodegeneration"

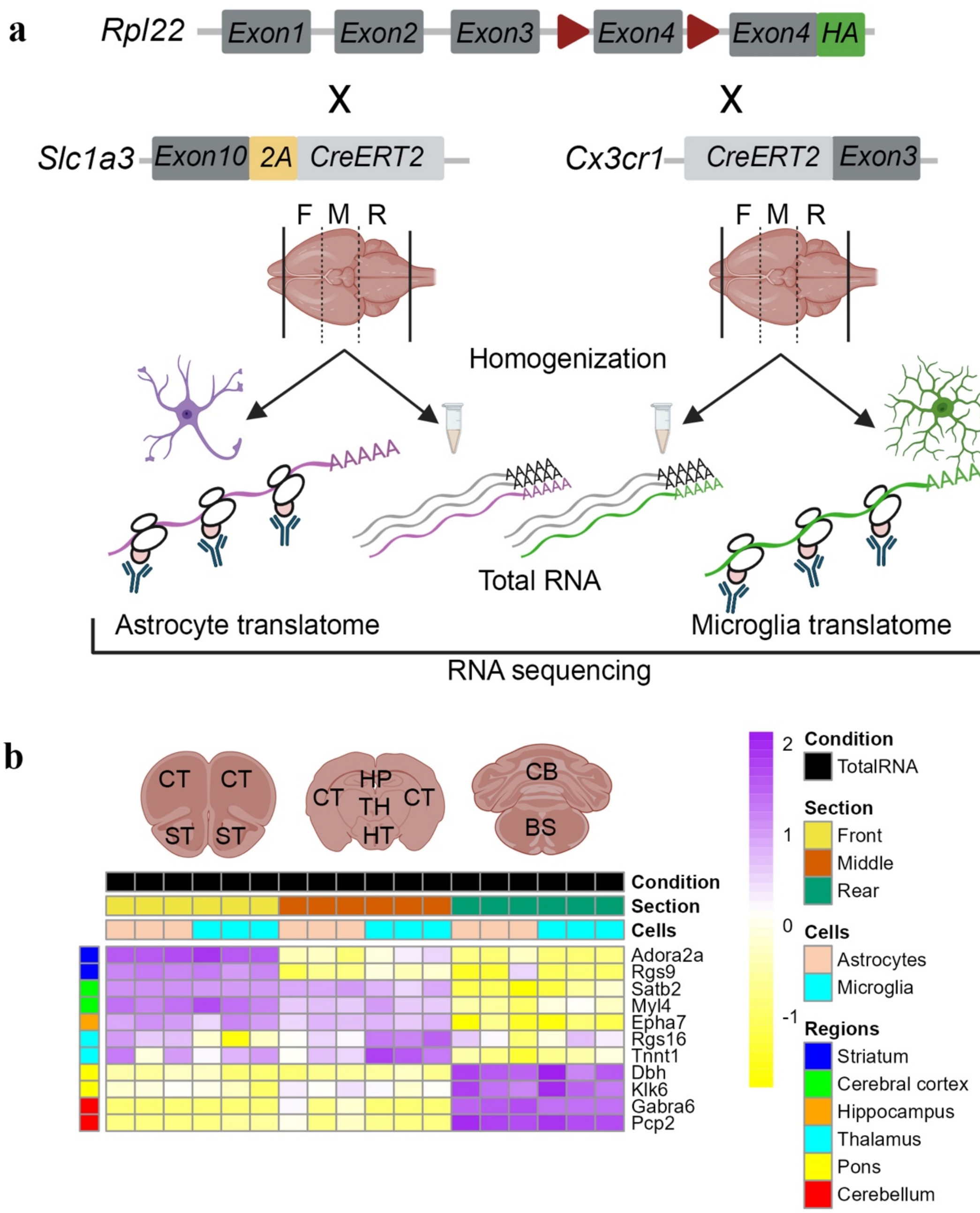


**Supplementary Figure 1. Strategy for cell type-specific translatome analysis across brain regions.** (a) Genetic approach for cell type-specific RiboTag expression. Schematic shows Rpl22-HA allele structure, cell type-specific Cre driver lines (Slc1a3-CreERT2 and Cx3cr1-CreERT2), and experimental workflow for isolating cell type-specific ribosome-associated RNA. Created with BioRender. (b) Validation of regional sampling and cell type specificity. Top: Anatomical reference showing sampled regions (CT: cortex, ST: striatum, HP: hippocampus, TH: thalamus, HT: hypothalamus, CB: cerebellum, BS: brainstem). Bottom: Heatmap of cell type-specific marker expression across conditions (TotalRNA vs RiboTag), brain regions, and cell types. Expression values are scaled from -2 (yellow) to 2 (purple); n = 3 independent samples per condition.

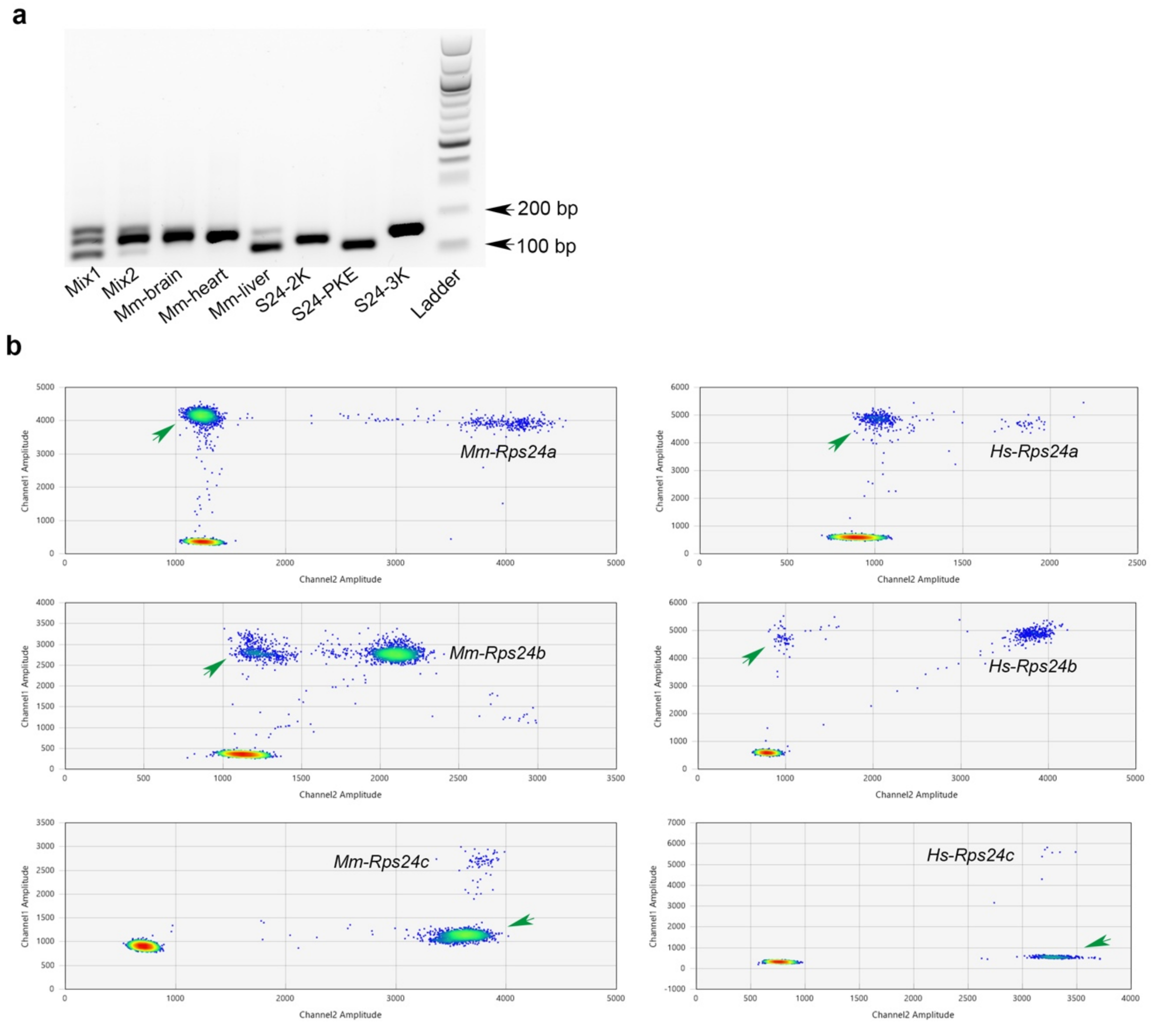


**Supplementary Figure 2. Technical validation of *Rps24* isoform detection.** (a) PCR validation of isoform-specific amplification. Agarose gel analysis of PCR products from *in vitro* RNA mix1 (equal concentrations of *Rps24a/b/c* isoforms), mix2 (modeling brain, 17% *Rps24a*, 80% *Rps24b* and 3% *Rps24c)*, mouse tissues, and unmixed isoform-specific controls of *in vitro* RNA (S24-2K, S24-PKE, and S24-3K). DNA ladder indicates 100 bp and 200 bp positions. Representative image from three independent experiments. (b) Droplet digital PCR optimization for isoform quantification. Two-dimensional amplitude plots showing distinct droplet populations for *Rps24/RPS24* variants in mouse (*Mm*) and human (*Hs*) samples. Green arrows indicate positive droplets; heat map shows droplet density (blue: low, red: high). Representative plots from n = 3 technical replicates per condition.

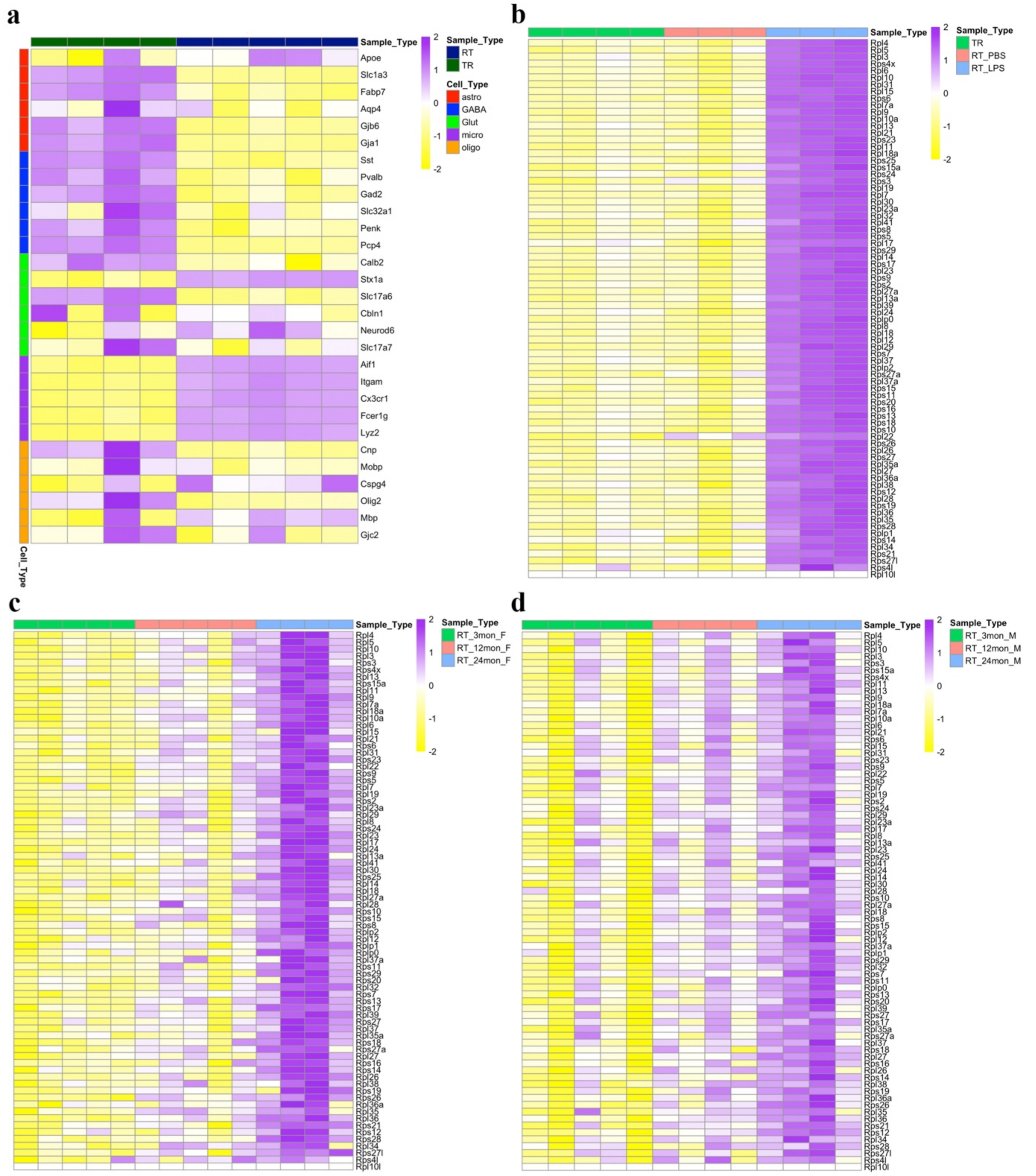


**Supplementary Figure 3. Analysis of cell type-specific markers and RP expression in inflammatory conditions.** (a) Cell type-specific marker expression in Kang dataset. Heatmap shows scaled expression values (-2 to 2) across neural cell populations. Sample types: TR (Total RNA, green), RT (RiboTag, blue). Cell types indicated by color bars: astrocytes (red), GABAergic neurons (blue), glutamatergic neurons (green), microglia (purple), oligodendrocytes (orange); n = 3 independent samples per condition. (b) RP transcript profiles in microglia under inflammatory conditions. Heatmap comparing RT_PBS (control) versus RT_LPS treatment. (c, d) Age-dependent changes in microglial RP expression in (c) female and (d) male. Samples from 3, 12, and 24 months shown. Expression values scaled from -2 (yellow) to 2 (purple); n = 3 mice per timepoint.

**Supplementary Discussion: *Rps24*/S24 isoform nomenclature**

Conventionally, gene and mRNA names use italics letters and protein names use non-italics letters. In humans, the names of genes and gene products are all uppercase whereas in mice only the first letter is uppercase. While trying to employ existing names for *Rps24*/S24 isoforms, we encountered numerous problems and were forced to devise a more systematic scheme. The original study of mouse *Rps24* that found the three main isoforms studied here used Roman numerals to label the 7 exons in order as, I through VII [36]. These transcripts containing 7, 6, or 5 exons were designated as S24b, S24a, and S24c, respectively (we named these as *Rps24a*, *Rps24b, and Rps24c*, respectively). A subsequent human study maintained this exon numbering but reported that exon V and isoform S24b (isoform *RPS24a* in our nomenclature) were absent from humans [37]. However, later work confirmed the presence of all three isoforms in both species[50]. In their labeling scheme [50], the exons labeled as V, VI, and VII by Xu et al. [36] were labeled by Gazda et al. [50] as VII, V, and VI, respectively. Furthermore, Gazda et al. [50] used numbers 1, 2, and 3 to refer to the isoforms called S24a, -c, and -b, respectively by Xu et al. [36]. More recently, Gupta and Warner [38] used Arabic numbers to label the exons in ascending order and used a transcript naming convention that progressively follows the number of exons excluded. More specifically, they named the transcript that excludes none of the exons as AA', the transcript that excludes 1 exon (exon 6) is called B, and the transcript excludes 2 exons (numbered 5 and 6) is called C. More recent publications have employed nomenclatures that further deviate from the original one used by Xu et al., [36]. For example, the terms small, large and extra-large have been used to distinguish transcripts that Xu et al. [36] called S24c, -a and -b, respectively [74]. While this is logical for the transcript isoforms, the protein isoform sizes have the reverse order, where translation of the shortest transcript makes the longest protein. The most recent nomenclature system is more complicated [42]. It starts with a different system for naming the exons. Those labeled IV, V, VI and VII by Xu et al. [36] (and 4, 5, 6, and 7 by us) were named 4, 18bp, 22bp and 6, respectively. This is then used to make transcript names ex4:ex6, ex4:22bp, and ex4:22bp/18bp that refer to Xu et al. [36] transcripts named S24c, -b and -a, respectively. Using the Xu et al. [36] numbering convention, the Park nomenclature is based on whether exon IV is spliced to exon VII (ex4:ex6) or to exon V (ex4:22 bp), and for transcripts in which exon V is spliced to exon IV, whether exon VI is also spliced in (ex4:22bp/18bp). Furthermore, databases like Ensembl number certain exons differently for each transcript, making discussions of specific exons cumbersome, and give different names (numbers) to the same transcript from mouse versus human. Importantly, a systematic approach for distinguishing the protein isoforms was essentially absent.

To address these inconsistencies, we developed a unified nomenclature system for both mouse and human variants that was logical and easy to remember. Therefore, like Gupta and Warner [38] we labeled the 7 exons in order with Arabic numerals 1 through 7. For transcript isoforms, we adapted the convention from Gupta and Warner [38] renaming AA', B, and C as *Rps24a, Rps24b, and Rps24c*, respectively. For the protein isoforms ending with KK or KKK, we find it useful for oral discussions to refer to them as S24-2K and S24-3K to constrain misunderstandings. Our system currently excludes the 3bp exon reported in humans [42], as it was not detected in our mouse transcriptome data.
